# Differential analysis of RNA-Seq incorporating quantification uncertainty

**DOI:** 10.1101/058164

**Authors:** Harold Pimentel, Bray Nicolas L., Suzette Puente, Páll Melsted, Lior Pachter

## Abstract

We describe a novel method for the differential analysis of RNA-Seq data that utilizes bootstrapping in conjunction with response error linear modeling to decouple biological variance from inferential variance. The method is implemented in an interactive shiny app called sleuth that utilizes kallisto quantifications and bootstraps for fast and accurate analysis of RNA-Seq experiments.

---

RNA-Seq technology has largely replaced microarray measurement as a tool for identifying gene expression differences in comparative and clinical analyses of RNA samples^1^. In analogy with microarrays, differential analysis of RNA-Seq experiments requires careful assessment of variability of gene expression from few replicate samples, so as to be able to identify biologically relevant expression differences between conditions^2,3^. However there are also key differences between the technologies. While microarrays measure cDNA hybridization intensities at predefined probes, RNA-Seq provides a *de novo* sampling of the transcriptome, a feature that makes it much more powerful for detecting transcription of individual isoforms of genes, but that also complicates differential analysis.

Many methods have been developed for differential analysis of RNA-Seq data^4^. Some of these seek to translate ideas developed for microarray analysis to the RNA-Seq setting^5^ whereas others are based on models tailored to RNA-Seq^6–10^. One of the key differences between RNA-Seq and microarray technology is that the data of the former consists of counts of reads rather than intensities measured at probes, and there has therefore been considerable effort devoted to exploring appropriate distributions for the modeling of count-based data in the RNA-Seq context^2,3,5,11–14^However the question of how to best utilize RNA-Seq data for differential analysis continues to be debated, with disagreements persisting on some of the most basic questions such as how to measure the abundance of genes^4^, whether there is sufficient power to test for differences in abundance of individual isoforms^15,16^ and how to best utilize biological replicates^17^.

Part of the reason for the continuing uncertainty regarding how best to analyze RNA-Seq data is the lack of agreed upon standards for testing and benchmarking methods. In most cases accuracy claims are based on simulations of counts of reads from distributions assumed in the models, rather than simulations of raw reads,^3,5,10,12,14,18,19^. Such count-based simulations typically discount the effects of ambiguously mapping reads and fail to capture both the possibilities for, and challenges of, isoform-specific differential analysis. Even when simulation studies are based on reads, they are sometimes restricted to a small portion of the transcriptome^20,21^ thereby biasing results due to the dependence of some methods on transcriptome-wide data for obtaining variance estimates from replicates. Studies utilizing biological data frequently make use of questionable choices for “ground truth”, e.g. utilizing results from microarrays, or from a differential analysis with a method that is imperfect^20^. The resulting benchmarks are therefore
difficult to interpret.

In this work we describe a novel approach to differential analysis of RNA-Seq data, a comprehensive framework for benchmarking our method and others that is unprecedented in its scale and scope, and interactive visualization software for exploring the results of our method and the data they are based on. The latter is crucial for providing transparency in assessing our results, and has the benefit of offering users a convenient tool for exploratory data analysis. Throughout the paper we use the name sleuth to refer both to our statistical method, as well as the app that allows for working with and exploring results.

The motivation for the conceptual approach underlying sleuth is illustrated in Figure 1. A key element of sleuth is borrowed from previous work^3,5^, namely shrinkage to stabilize variance estimates from few samples. But sleuth is able to leverage recent advances in quantification^22^ to obtain error estimates for quantifications that can in turn be used to decouple biological variance from inferential variance before shrinkage. A few other methods have attempted to compute and utilize error estimates on quantifications, but some major computational and statistical hurdles have been difficult to overcome. For example, the BitSeq method^10^ obtains quantification error estimates via Markov Chain Monte Carlo sampling, a process that requires significant, and in some cases prohibitive, computational resources. An update to the initial paper introduces quantification using variational Bayes, however it is not recommended for use in differential analysis^23^. The Cuffdiff 2 method^24^ also performs sampling to assess variance arising from quantification but has some model limitations: Cuffdiff 2 fails to accurately assess the increase in variance due to reads mapping ambiguously across the genome. EBSeq^15^ also models inferential variance, but does so in discrete classes that only act as a proxy for high inferential variance.

Thus, sleuth is able to improve on traditional “count-based” methods by utilizing improved estimates of transcript and gene abundances in a flexible and powerful statistical framework. The sleuth concept is illustrated by example in Figure 1 via the tracing of genes whose abundances are difficult to estimate. In the example shown, which is based on a gene-level analysis of data from Bottomly et al. ^25^, genes with high inferential variance (red dots in Figure 1a) are lumped together with genes with high biological variance by DESeq2 (Figure 1b). This makes it difficult to correctly assign a high total variance to those genes prior to differential analysis (Figure 1c). Unlike DESeq2, by decoupling biological and inferential variance (Figure 1c, d), sleuth assigns a high total variance to most of the genes with high inferential variance (Figure 1e).

The key innovation in sleuth, namely the explicit modeling of biological and inferential variance, is performed with a response error model (see Methods and Supplementary Materials). The model is simple, transparent and its parameters are easily interpretable. Inference of parameters is straightforward (see Supplementary Materials). Thus, when coupled with kallisto^22^, which is fast in both the quantification and variance estimation, sleuth provides a statistically rigorous, flexible and efficient solution for RNA-Seq analysis.

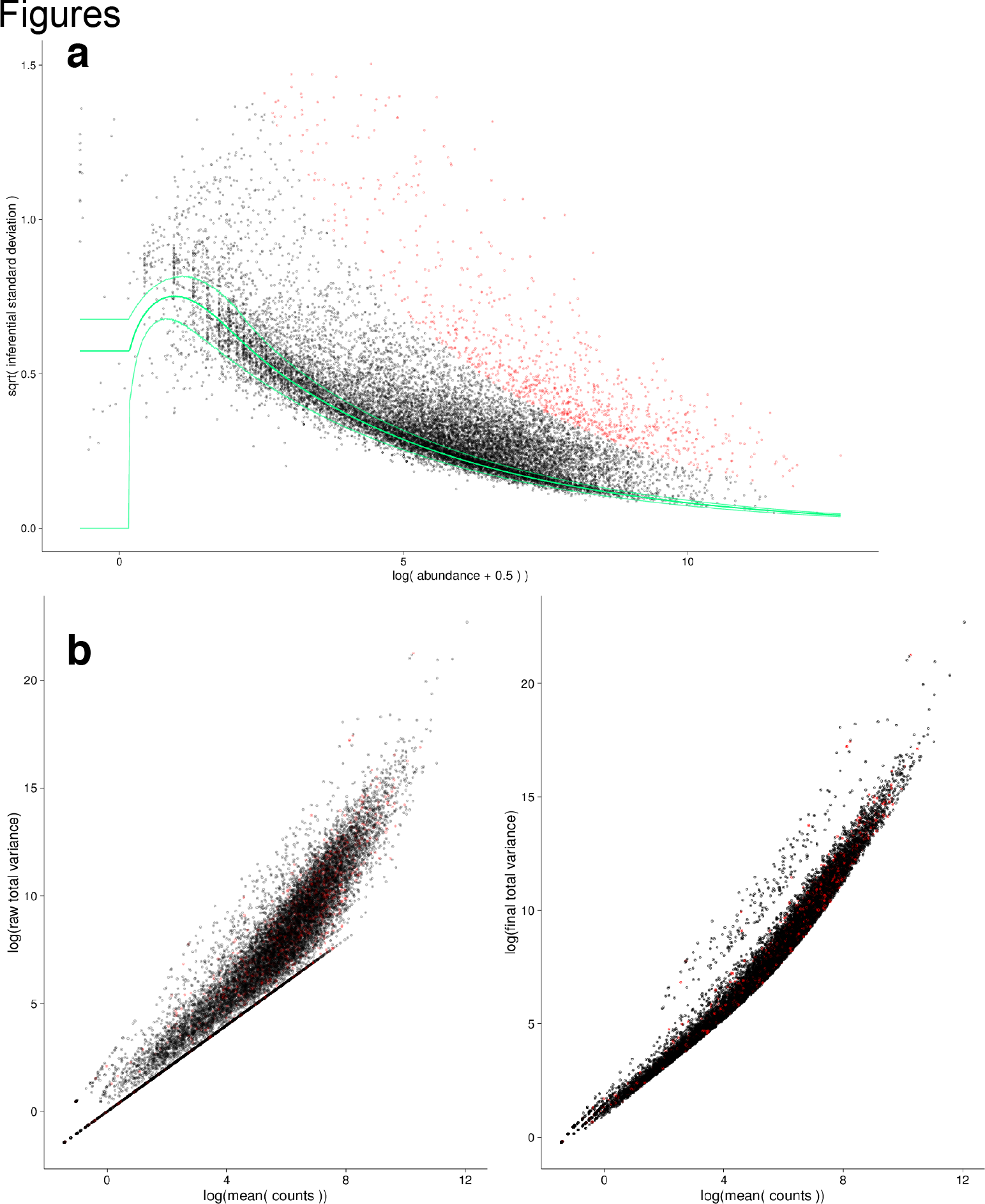

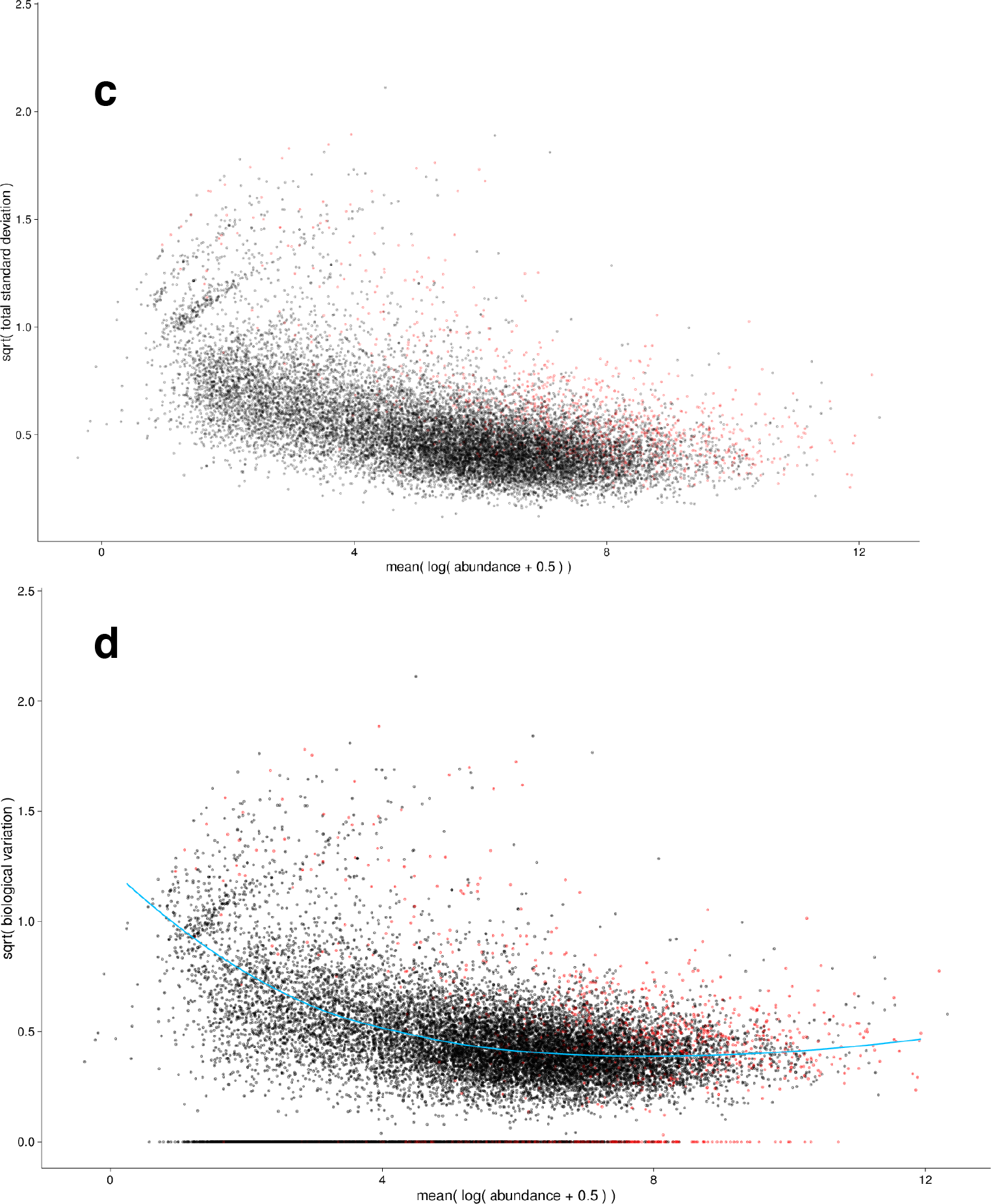

**Figure 1:**
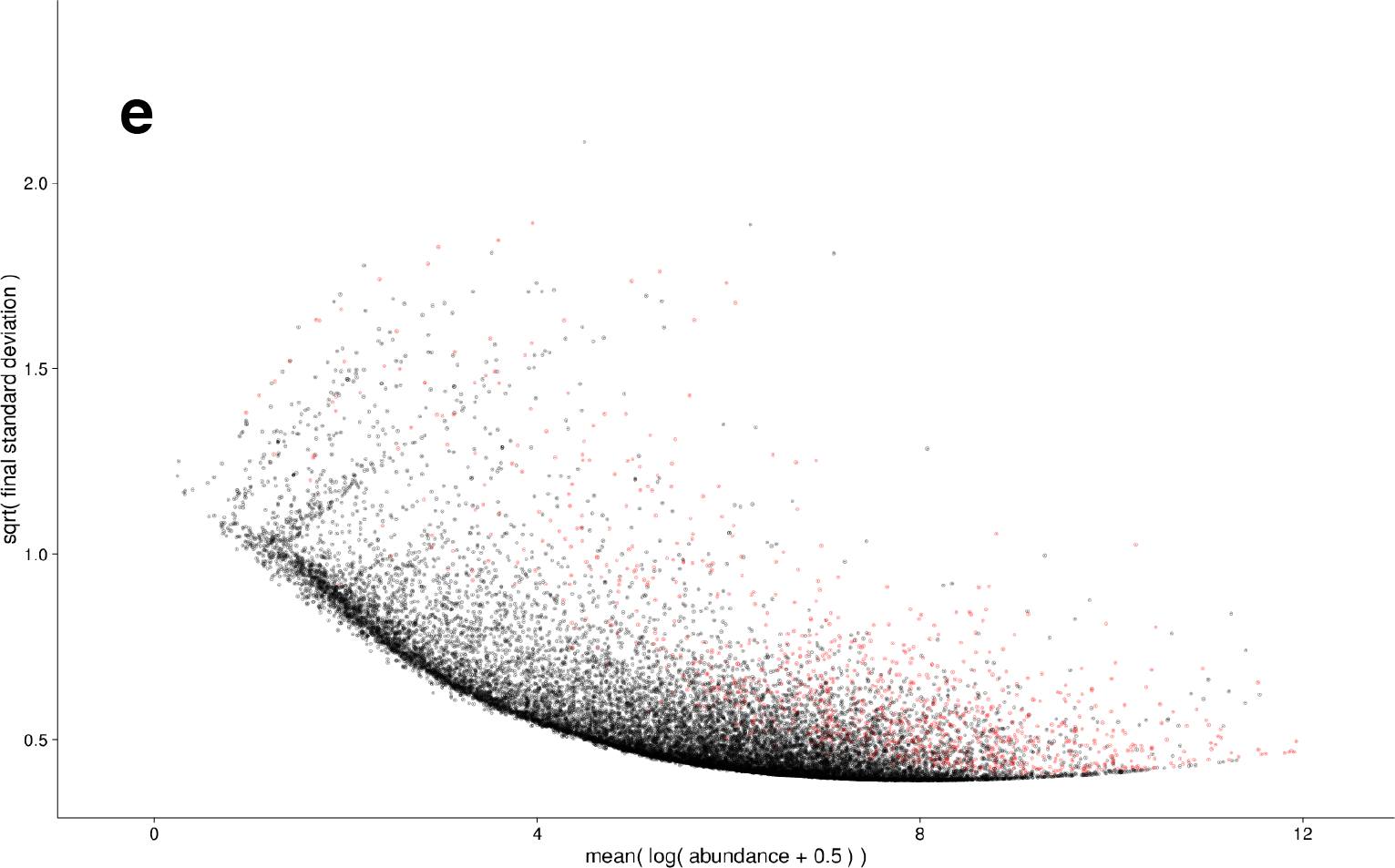
The effect of modeling inferential variance in sleuth at the gene level. Outliers are colored red and traced across all plots. A point is an outlier if the variance is greater than 100 times the interquartile range plus the upper quartile. (a) Inferential variance on sample **XXX**. The x-axis is the gene abundance, and the y-axis is the bootstrap estimate of the inferential variance. The blue lines represent the 5% confidence bound, mean, and 95% confidence bound expected under the Poisson model. (b) Mean expression versus total variance as estimated by DESeq2. The left panel contains the raw estimates of the variance. The right panel contains the smoothed estimate of the variance. Note that the outliers are fairly randomly distributed across variance and expression patterns. (c) Raw total variance as estimated by sleuth. Again, the outliers are fairly randomly distributed since they do not consider the inferential variance. (d) Biological variance as estimated by sleuth once the inferential variance has been removed. The blue line represents the mean-biological variance relationship modeled in sleuth. Note that in many cases the inferential variance is greater than the biological variance resulting in an estimate of biological variance equal to zero. (e) Final total variance as modeled by sleuth. Note that almost all of the outliers have higher abundance than the nonoutliers due to high inferential variance.

## Results

### Modeling inferential variability

The fundamental issue in differential analysis of RNA-Seq experiments is quantifying the variance in experiments so that true differences in expression can be identified as such. There are multiple sources of variance that contribute to the total variance observed between samples in an RNA-Seq experiment which we group into two classes: (1) “biological variance” which is a term used to describe variance in transcript abundance of biological and experimental origin and (2) “inferential variance” which is a term we use to describe both variance in the number of reads sequenced from a transcript due to the random nature of sequencing as well as the variance that emerges as a result of the statistical nature of transcript abundance estimation with ambiguously mapping reads. In the absence of ambiguously mapping reads there is no increase in inferential variance as the origin of each read can be inferred exactly. However when reads map to many transcripts the read counts must be deconvoluted to obtain the abundance estimates^8^ leading to uncertainty in abundance estimates which translates into variance in quantification across samples.

Response error modeling^26^ allows for the separate modeling of biological variance and inferential variance. The use of the bootstrap during quantification allows us to estimate the inferential variance directly for each sample^22^, whereas biological replicates allow us to estimate the total variance, albeit via shrinkage due to the limited number of replicates in most experiments (see Methods). The biological variance can then be estimated (see Supplementary Materials). In Figure 1 we illustrate this procedure by first showing the inferential variance versus the mean at the gene level (Figure 1a). Notice that for many genes the inferential variance is much higher than the expected Poisson variance (which is assumed by most methods). We track the outliers in red throughout all of Figure 1. Figure 1b shows the total variance when estimated using DESeq2 (raw estimate on left, final shrinkage estimate on right). The inferential variance outliers are randomly distributed throughout the mean variance relationship since there is no additional information about the inferential variance. Figure 1c shows the raw estimate of the total variance in sleuth without any knowledge of the inferential variance. Here, the red points are randomly distributed, while Figure 1d shows the biological variance after the inferential variance has been removed. Many of the red points are distributed near biological variance zero. These points have more inferential variance than biological variance. The points with nonzero biological variance are a mix of points where the total variance is very high or inferential variance is very low. Sleuth performs shrinkage on these estimates (blue line). Figure 1d displays the final estimates of the total variance, which are the smoothed biological estimates plus the inferential variability estimates (Figure 1a). Due to the shrinkage procedure, the red points have the highest total variance and are penalized heavily as a result unlike the final variance in Figure 1b and 1c.

### Improved accuracy in differential gene analysis

In order to test the performance of sleuth in differential gene analysis we examined both simulated and real data and compared it to numerous other widely used methods. Our simulation was derived from an experiment with two conditions and three replicates in each condition (see Methods). We simulated biological variance (dispersion) according to the negative binomial model for counts used by DESeq2^19^ (see Methods). To be able to accurately assess performance, each simulation was performed with 20 replicates.

Figure 2 shows the result of our method in identifying differentially expressed genes, alongside results from Cuffdiff 2^24^, DESeq^3^, DESeq2^19^, EBSeq^15^, edgeR^11,18^, voom^5^, and log fold change^20^. The sensitivity of sleuth is higher than all other methods in the false discovery rate (FDR) range of usual interest and beyond, up to FDR 0.3. The figure also shows that as expected, DESeq2 has more power than DESeq at all the relevant FDRs, and that the naïve approach of ranking genes by log-fold change produces poor results. Even when benchmarked in simulation conditions favorable to traditional “count-based” methods, sleuth outperforms other programs (Supplementary Figures 3-8). We also examined the effect of different filtering strategies on performance by comparing sleuth with other programs on a common filtered set of genes, showing that sleuth maintains its advantage independent of filtering (Supplementary Figures 914).

**Figure 2:**
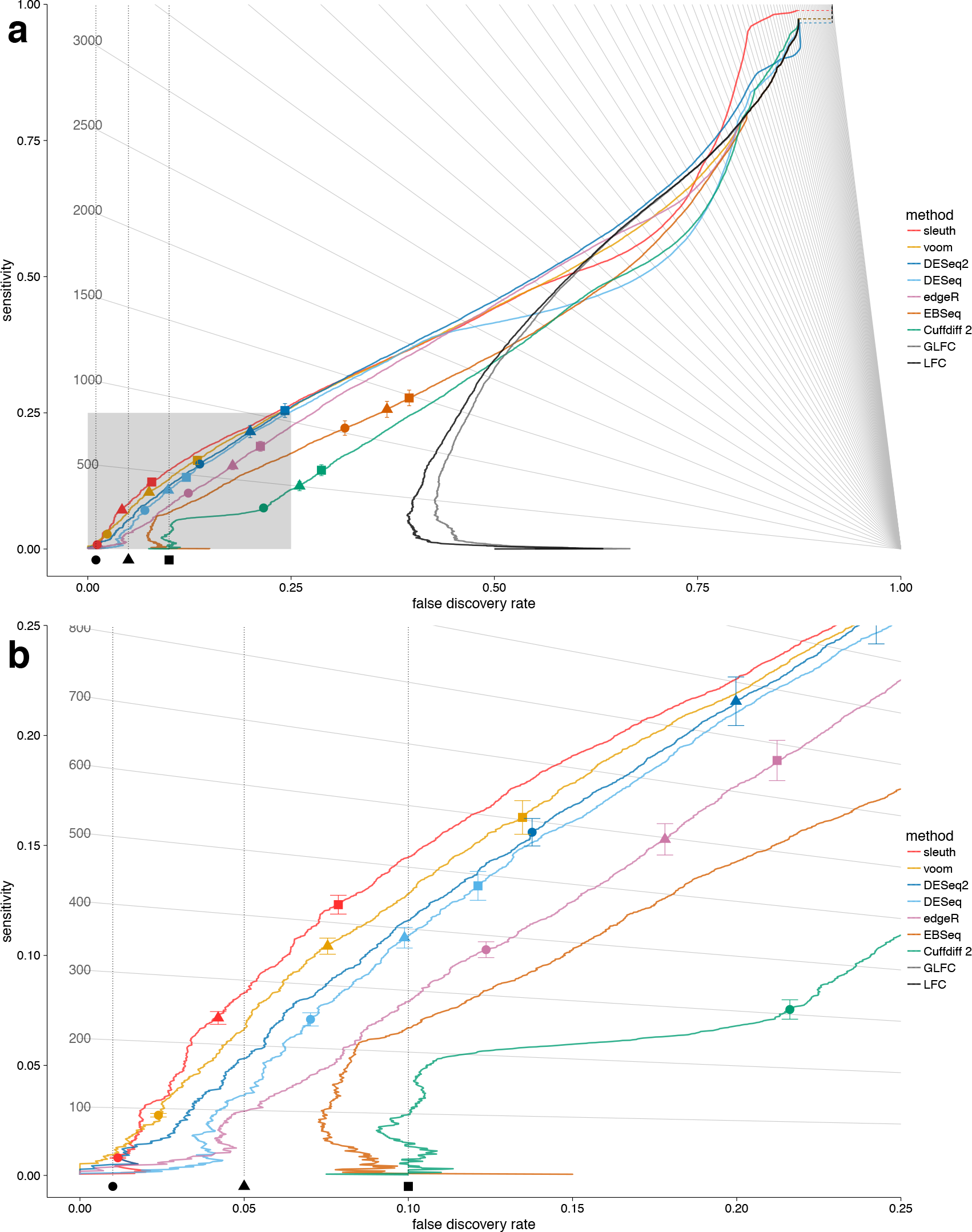
Sensitivity versus FDR in the effect from experiment simulation at the gene level. The x-and y-axes show the true false discovery rate and sensitivity, respectively, and for each program one has a curve showing how those values change as one moves down its ranking of all genes passing its filter. The circle, triangle, and square on each curve show where in its ranking that program estimates an FDR of 0.01, 0.05, and 0.10 respectively. Ideally, each symbol would lie directly above the corresponding symbol on the x-axis indicating a true FDR of 0.01, 0.05, or 0.10. Also displayed are isolines for a constant number of genes being called differentially expressed. Where an isoline intersects with the curve for a given program shows its performance when looking at that many genes from the top of its ranking. The FDR lines were averaged over 20 replications of the simulation. Panel (a) shows the entire plot while panel (b) is zoomed into the region where true FDR is less than 0.25.

### Estimation of false discovery rate

Since the control of the false discovery rate is fundamental for identifying differentially expressed genes in experiments with few replicates, we examined carefully the accuracy of methods in self-reporting their false discovery rates^27^. This was easy to do with our simulated data where the truth was known. In Figures 2 and figure 4 the circles, squares, and diamonds represent the average estimated FDR output by each program across the 20 replicates performed for each simulation scenario. Other than sleuth and voom, other methods significantly underestimated the FDR with several methods reporting an estimated FDR of 0.01 when the true FDR was greater than 0.1. While sleuth overestimates the FDR, this error is conservative, i.e. fewer genes are reported, yet they are highly enriched for being differentially expressed.

To test whether our results translated to FDR estimation accuracy on real data, we repeated an experiment from the DESeq2 paper^19^. Using the Bottomly data set^25^, which contains multiple replicates from two mice strains (10 and 11 respectively), we created a training set by randomly selecting 3 versus 3 samples and using differential expression results from the remaining 7 versus 8 as the “truth”. Each method was compared to itself to see how well it could recapitulate its results with a smaller set of data and how it controlled the FDR as assessed by comparing to the results of the high replicate analysis. We iterated this procedure 20 times. Figure 3a shows that as in the simulation, sleuth and voom are the only methods able to estimate their FDR to within a reasonable approximation.

To see whether the consistency experiment provided results concordant with those of our simulations, we performed the consistency experiment with simulated data. The results are shown in Figure 3b, which illustrates both that our simulated data recapitulates the results on real data, and that the self-referential FDRs are good proxies for true FDRs, thus validating the reliability of the DESeq2 consistency experiment.

**Figure 3:**
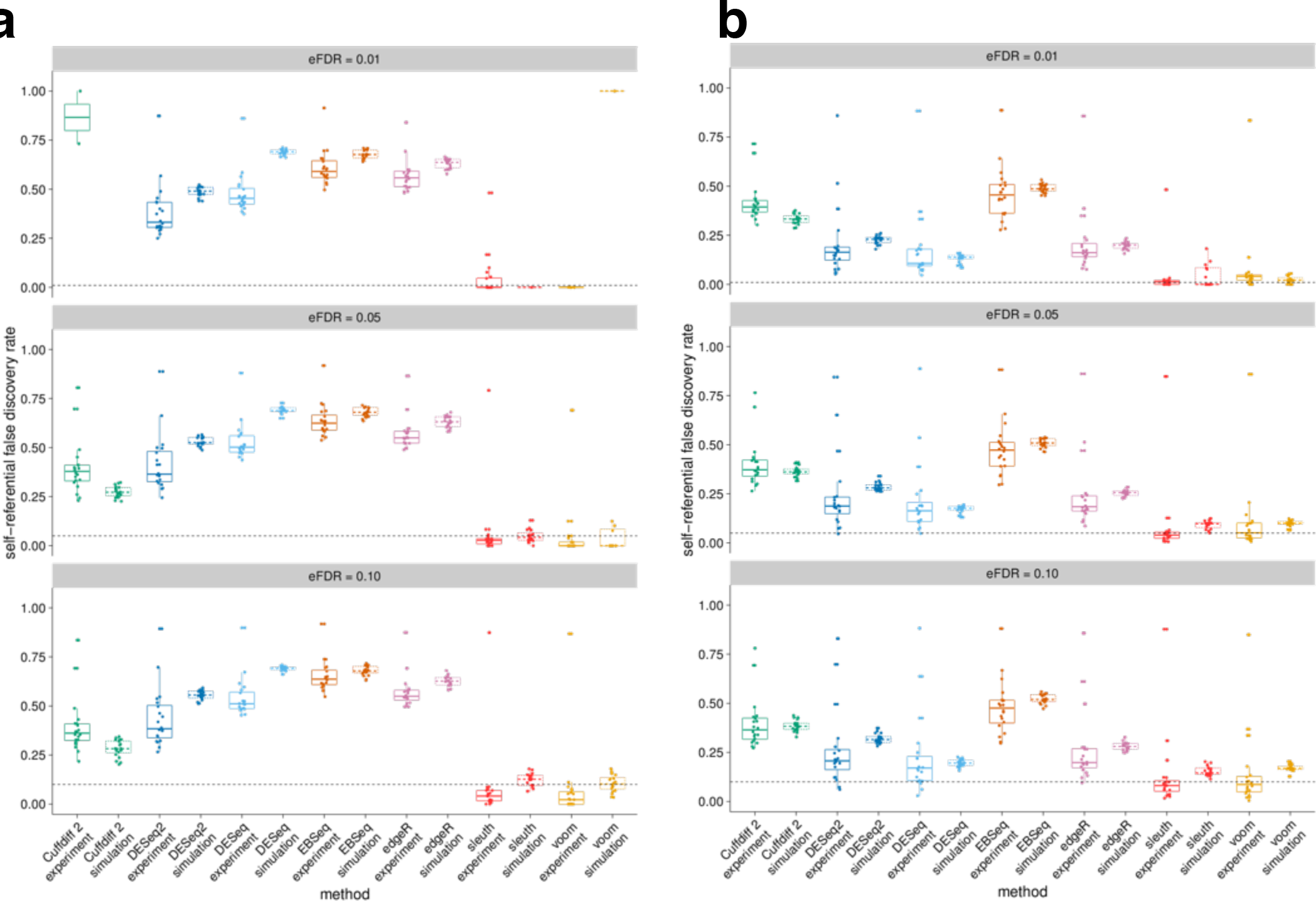
Self-referential FDR for the Bottomly data set and our simulation at the (a) isoform level, and the (b) gene level. The suffix “experiment” refers to the Bottomly data set whereas “simulation” refers to our simulated experiments to mimic the Bottomly resampling experiment. The panels from top to bottom display the true FDR for each program when it estimates the FDR as 0.01, 0.05, and 0.10, respectively. The dashed horizontal line represents the target FDR. Only sleuth and voom seem to control the self-referential FDR reasonably well at both the isoform and gene level.

### sleuth allows for isoform-level differential analysis

While RNA-Seq has become the standard technology for *gene*-level differential analysis, there has been some debate about its suitability and power for *isoform*-level differential analysis. In previous work, we and others have provided examples of how isoform-level differential analysis can highlight interesting splicing and differential promoter usage between conditions^24,28^ but there has been debate about the significance and reliability of such results^16,20^.

In order to examine this question, we repeated the gene-level analysis at the transcript level (see Figure 4). We confirm previous findings that because increased testing is required for isoform-level analysis, there is a decrease in sensitivity in comparison to gene level analysis. However we also find that sleuth can still control the false discovery rate at the isoform level while calling many isoforms differentially expressed. Interestingly, while there is less power to discover differentially expressed isoforms, our simulations show that at a given FDR the number of differentially expressed features is fairly similar to that of genes (isolines in Figures 2,4, and Supplementary Section 9). Moreover, when simulating from a scenario in which isoform abundances change independently between conditions (Supplementary Figures 1-4), we find highly significant improvements in sleuth with respect to other methods. The same is also true for the correlated effect simulation. In addition, we tested BitSeq^10^ (Supplementary Figures 15 and 16) on a single sample as its run-time was prohibitive on the entire simulation set. We found BitSeq performed well overall although sleuth outperformed it when the true FDR was less than 0.12.

**Figure 4:**
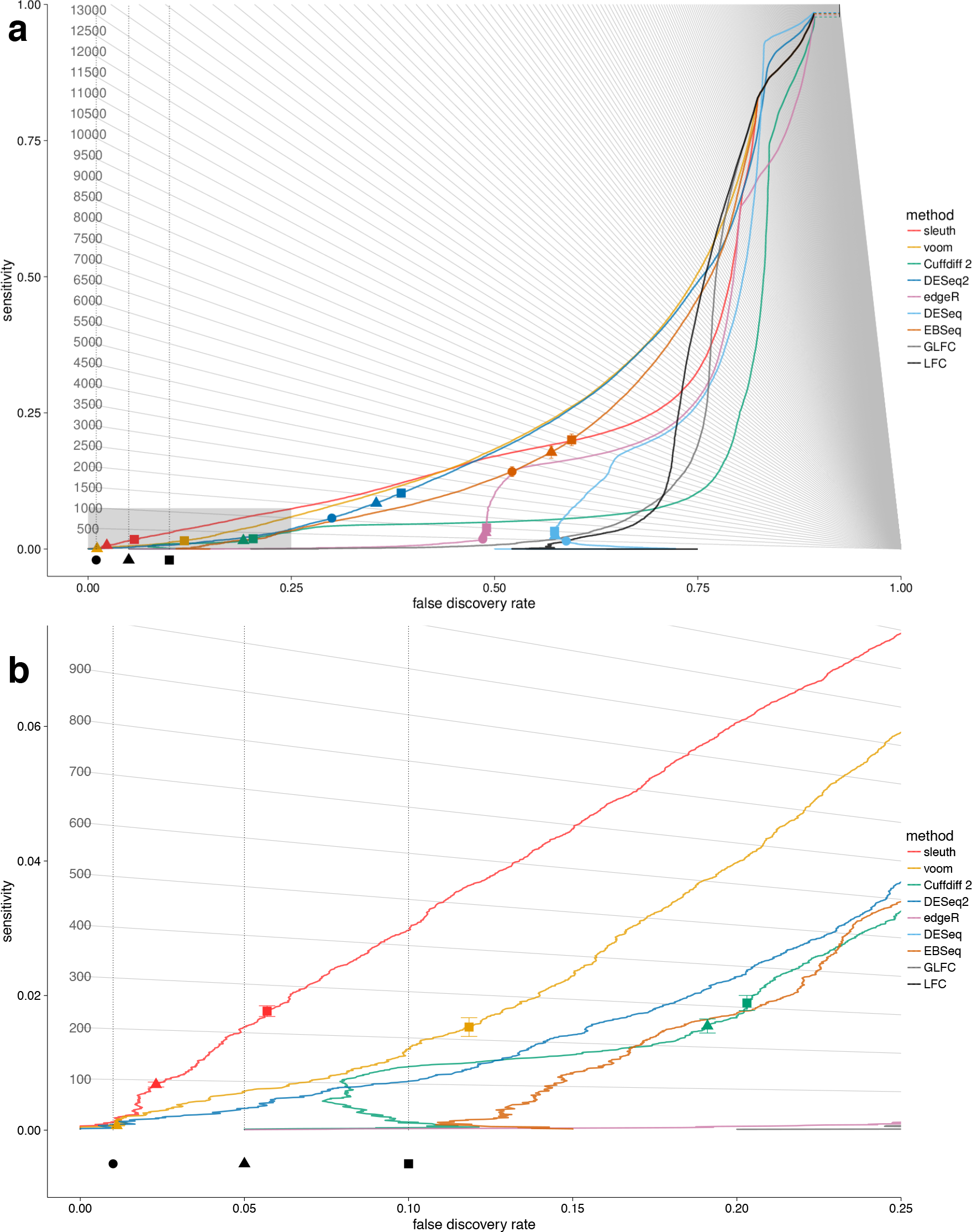
Sensitivity versus FDR in the effect from experiment simulation as in Figure 2 at the transcript rather level.

### Interactive exploratory data analysis using sleuth

The interpretation and analysis of RNA-Seq data is complicated by a number of factors: the large quantity of reads sequenced in typical experiments (many millions) and the large number of transcripts / genes (typically tens of thousands) make it difficult to interactively examine the data. However exploratory data analysis is important both for understanding how to analyze the data and in the formation of hypotheses about the results. To address this issue, and to make it possible to evaluate and assess the results of sleuth, we have developed a Shiny^29^-based interactive app for examining sleuth results.

Figure 5 shows some screenshots from a sleuth analysis of the Bottomly data^25^. Figure 5a shows the principal component analysis of the data set colored by the different conditions. One can see that the first two principal components do not segregate the data by experimental condition (mouse strain). Figure 5b shows how one can use the drop-down menu to change the coloring, revealing that the first two principal components seem to explain some of the variation due to the batch. In addition, there are many other features assisting in exploring the data, such as the ability to view, sort and search the table of differential expression results. For example, sorting by the inferential variability and then by largest p-values, we find transcript ENSMUST00000113388, which is not reported as differentially expressed by sleuth, but is reported as differentially expressed by both voom and DESeq2. This is likely due to the high inferential variability which is not being properly assessed and adjusted for by those programs. The transcript name can be pasted into the “transcript view” window and the distribution of inferential variability can be explored with boxplots describing the variability within each sample (Figure 5c).

**Figure 5:**
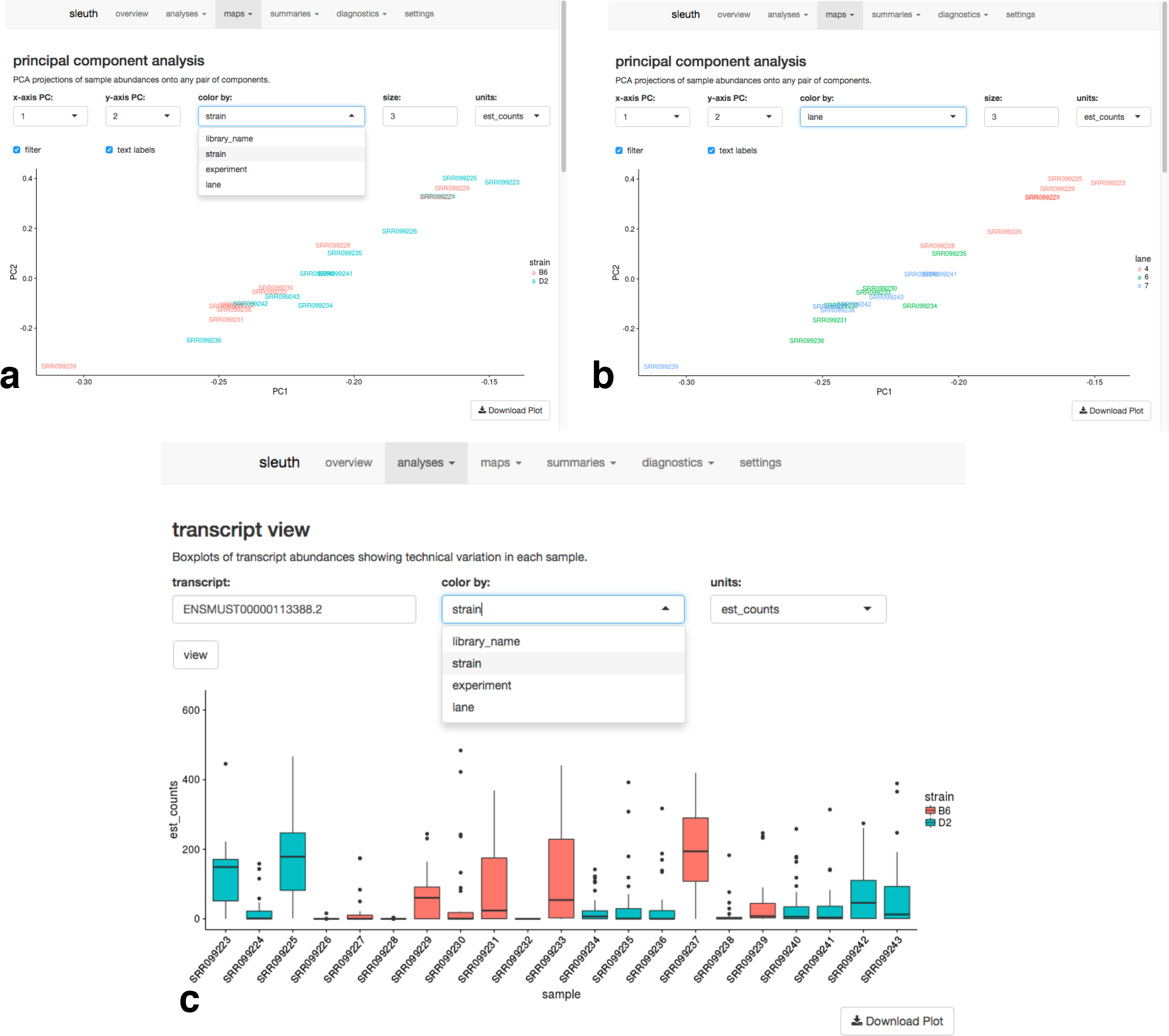
Interactive sleuth live Shiny interface on complete Bottomly data set. (a) PCA plot colored by strain shows that the strain does not explain much of the variance in the first two principal components. The coloring can be changed immediately by drop-down as shown in (b) which indicates that there are possible lane effects. (c) Sample specific bootstraps for a particular transcript ENSMUST00000113388, which does not show differential expression by sleuth, but shows differential expression by limma and DESeq2. A possible explanation for this is that the inferential variance is quite high.

## Discussion

RNA-Seq experiments produce data that is complex in structure and rich in information. This data presents an unprecedented opportunity for studying transcriptional mechanisms and, but its analysis is fraught with challenges. In the case of differential expression, the large volumes of data and the large sizes of transcriptomes have made it difficult to explore results “by hand” in order to gain intuition and insight into the experiments. “Count-based” methods for differential analysis have been popular partly because they are simple in their approach and present researchers with numbers to examine that are easy to relate to. However the simplicity of count-based methods comes at a cost: by ignoring the complexity of ambiguously mapped reads they introduce biases that can have detrimental effects on results^24^.

The sleuth model provides a solution to a perplexing difficulty in RNA-Seq analysis: it offers a simple yet powerful framework for “counting” even when reads cannot be assigned unambiguously to transcripts, and therefore allows for robust and accurate RNA-Seq analyses. Our results show that by virtue of appropriately accounting for uncertainty in quantifications, sleuth is more accurate than previous approaches at both the gene and isoform levels. Crucially, the estimated FDRs reported by sleuth reflect the true FDRs, and therefore make the predictions of sleuth reliable and useful in practice.

The sleuth workflow has been deliberately designed to be simple so that it is interpretable and fast. The model was chosen in part because of its tractability and the Shiny framework for visualization was chosen for its portability. The modularity of the algorithm also makes it easy to explore improvements and extensions, such as analysis of more general transcript groups (e.g. as defined by shared exons, or 5’/3’ UTRs) and different shrinkage and normalization schemes to improve performance. As a result, when coupled with kallisto, which has dramatically reduced running times for quantification based on the idea of pseudoalignment, sleuth is a quick, accurate, and versatile tool for the analysis of RNA-Seq data.

## Online methods

### Model

We consider an additive response error model^26^ in which the total between-sample variability has two additive components: “biological variance” that arises from differences in expression between samples as well as variability due to library preparation, and “inferential variance” which includes differences arising from computational inference procedures in addition to measurement “shot noise” arising from random sequencing of fragments. The model is an extension of the general linear model where the total error has two additive components. Given a design matrix *x* we assume a general linear model for the (unknown) abundance **Y**_ti_ of transcript *t* in sample *i* in terms of fixed effects parameters beta and “noise” epsilon 
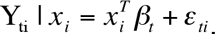

While the **Y**_ti_ are not directly observed, (normal) perturbations *ζ*_*ti*_ of them constitute the observed random variables **D**_ti_:

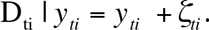

With some further assumptions one can derive that the D_ti_ are normally distributed (see Supplementary Materials) and the variance, which is the key to differential analysis, can be interpreted as the sum of biological (*ε*_*ti*_) and inferential (*ζ*_*ti*_) variance.

## Testing for differential expression

In comparing samples to identify differential expressed genes or transcripts, sleuth applies the likelihood ratio test where the full model contains labels for the samples and the reduced model ignores labels. Underlying the test is an estimate of the variances ***V***(***D***_*ti*_) where *t* ranges over the transcripts and *i* over the samples. The estimate for ***V***(***D***_*ti*_) used in sleuth is

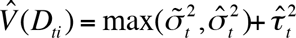
where 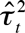 is the estimate of the inferential variance, 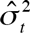 the raw biological variance and 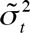 a shrinkage-based estimator of the biological variance. For details of how the individual variance estimates were obtained see the Supplementary Methods.

## Simulations

A null distribution for transcript abundances was learned from the largest homogeneous population in the GEUVADIS data set:59 samples of Finnish females^30^. We estimated transcript-level abundances with kallisto, then estimated parameters for negative binomial distributions (using the Cox-Reid dispersion estimator) to model count distributions using DESeq2.

After the null distribution was constructed, expression features (isoforms or genes depending on the type of simulation) were chosen to be differentially expressed. Transcripts with less than 5 estimated counts on average across the GEUVADIS samples were marked as too rare to be simulated as differentially expressed. A gene was assumed to pass the filter if at least one of its constituent transcripts passed the filter. In each simulation, 20% of the features that passed the filter were chosen to be differentially expressed at random. If the simulation had unequal size factors, random size factors were chosen from the set {1/3, 1, 3} such that the geometric mean equaled 1 similar to the simulation procedure in DESeq2. However, unlike the DESeq2 simulation procedure our size factors were chosen at random. Counts were generated from the negative binomial distribution after which reads were simulated using the RSEM simulator^7^.This resulted in about 30 million 75 base-pair paired-end reads per sample for a total of 13.8 billion reads overall (Supplementary Tables 2-5). Three types of simulations were performed:

### Independent effect simulation

Isoforms were chosen to be differentially expressed at random. This simulation was similar to the default in polyester^31^, however our simulation was performed across the entire transcriptome rather than just a few chromosomes. The simulations were generated with equal size factors. Effect sizes were chosen from a truncated normal distribution such that the minimum absolute fold change for differential transcripts or genes was 1.5.

### Correlated effect simulation

Genes (instead of isoforms) were randomly chosen to be differentially expressed. A direction (sign) for each effect size was chosen at random, then all the effects were simulated from a truncated normal with minimum absolute fold change 1.5. The simulation used random unequal size factors generated as described above.

### Effect from experiment

To mimic the types of changes seen in real experiments, fold changes were learned from Trapnell *et al.^24^* from the set of transcripts that either DESeq2 or sleuth found to be differentially expressed at FDR 0.05. Genes were chosen at random to be differentially expressed. The null mean counts were used to determine the rank of each transcript relative to its parent gene. These ranks were matched between the Trapnell data set and the null distribution learned from the GEUVADIS data set.

### Self-consistency experiment

In order to validate whether methods would produce similar results with less data, we performed an experiment similar to Love *et al.*^19^. For each iteration we randomly selected 3 samples from condition C57BL/6J and 3 samples from condition DBA/2J and ran each tool. The remaining samples were used as the “truth” by calling differentially expressed genes or transcripts using them. For each FDR level (0.01, 0.05, 0.10), we compared the results from the smaller data set to the larger data set for each tool. The FDR was then computed and plotted in Figure 4.

## Software notes

The following R programs were used to compile the results: sleuth 0.28.1, BitSeq 1.16.0,DESeq 0. 24.0. DESeq2 1.12.0, EBSeq 1.12.0, edgeR 3.14.0, limma-voom 3.28.2. When testing programs at the isoform-level, kallisto 0.42.4 was used to obtain quantifications. Cuffdiff 2.21 was used with alignments from HISAT2 2.0.1^32^. Subread (featureCounts) 1.5.0^33^ was used with alignments from HISAT2 to get raw gene counts. BitSeq was provided alignments from Bowtie 1.1.2^34^. All analyses in the paper are fully reproducible through the Snakemake system^35^.

## Acknowledgements

HP and LP were partially supported by NIH R01 DK094699 and NIH R01 HG006129. We thank Daniel Li, Alex Tseng, Pascal Sturmfels for help with implementing some of the interactive features in sleuth.

